## Supplementary figures and images for "The dual role of Spn-E in supporting heterotypic ping-pong piRNA amplification in silkworms"

### Fig. EV1

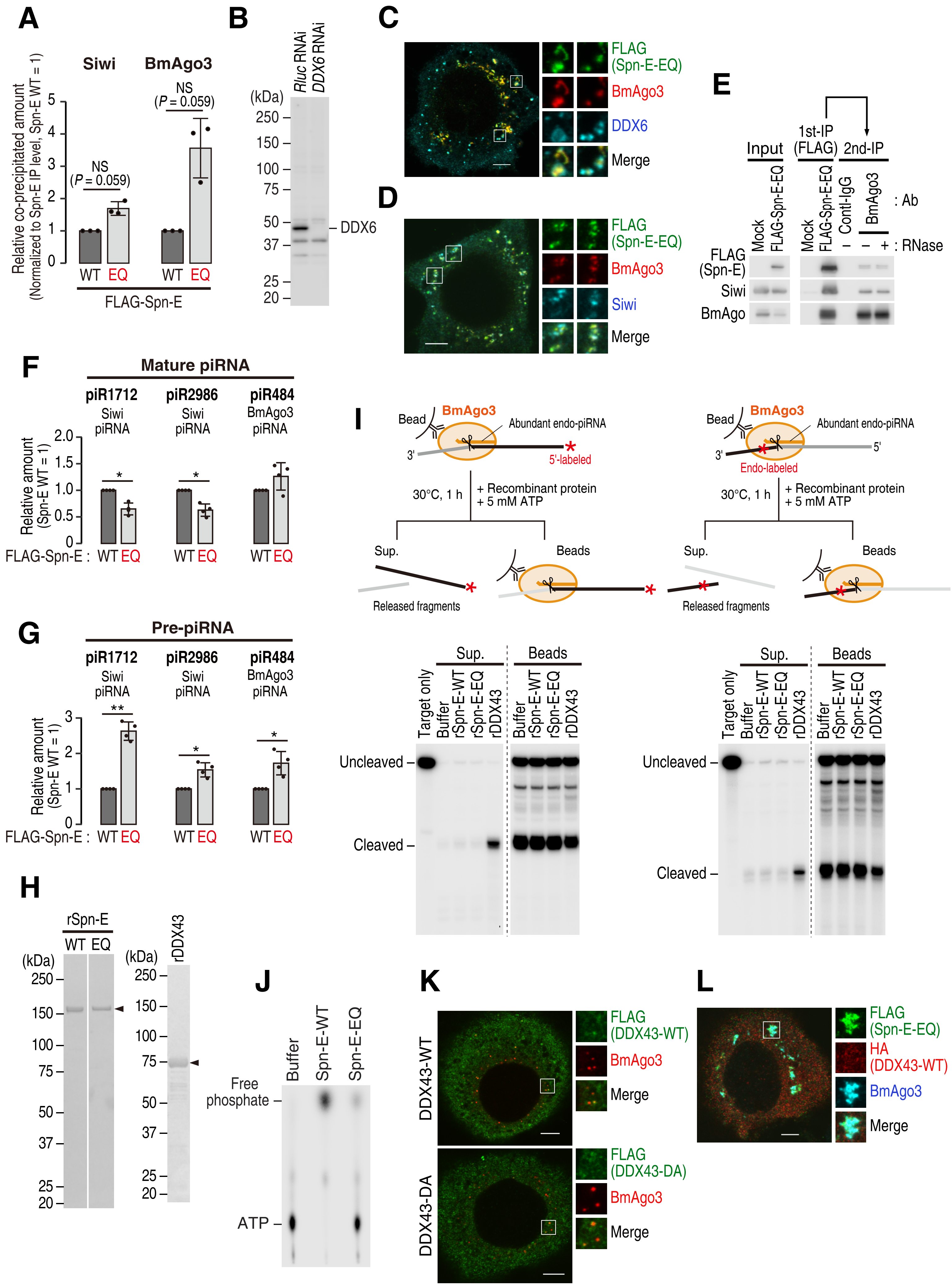

### Fig. EV2

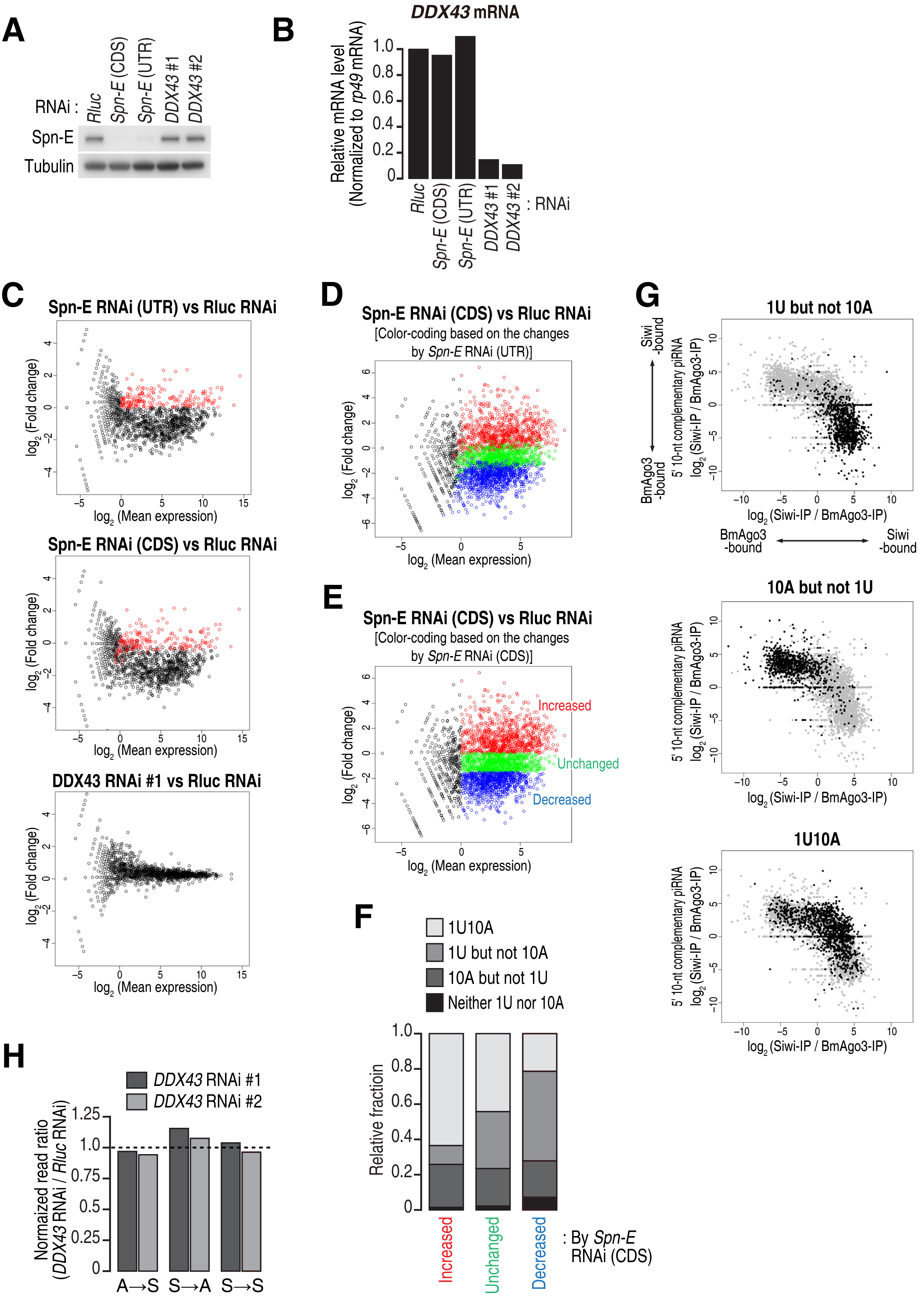

### Fig. EV3

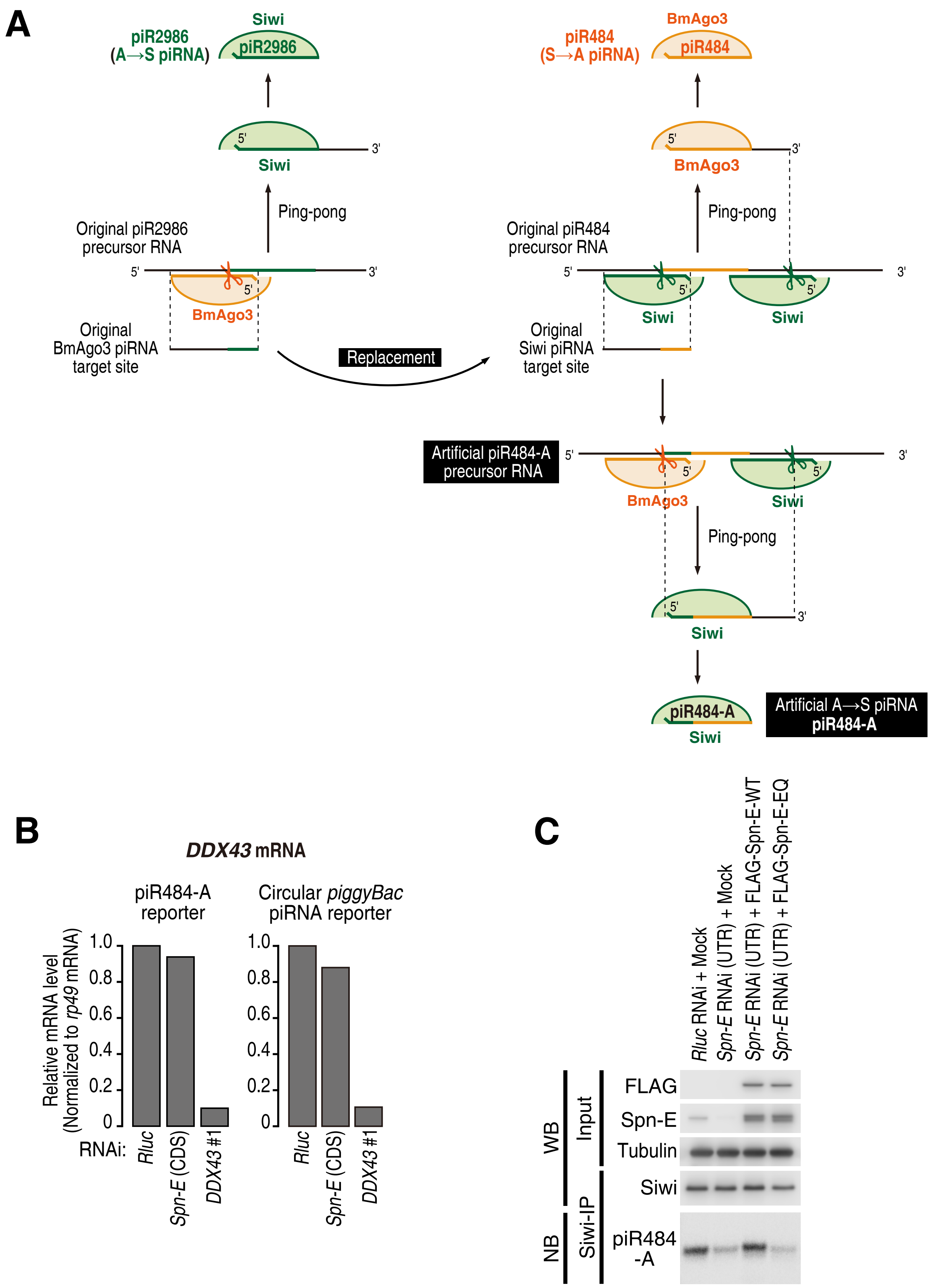
