## Supplementary material for "The dual role of Spn-E in supporting heterotypic ping-pong piRNA amplification in silkworms": Table EV1

**Table EV1. Oligonucleotide sequences used in this study.**

| Plasmid construction | 5′- 3′ |
|---|---|
| Spn-E-E251Q-F | TCTTGATCAAGTTCACGAAAGAGGACAAGAGATG |
| Spn-E-E251Q-R | GTGAACTTGATCAAGAATAACATGGGTGTATTC |
| FLAG-DDX43-F | ATGACGACGATAAGACTGATCGCGACGATGATGA |
| pIEx1-DDX43-R | CTGAGGTTAATCACTTACCATCTGCCTCTCCCTCTAC |
| FLAG-R | CTTATCGTCGTCATCCTTGTAATC |
| pIEx-3'out-F | GTGATTAACCTCAGGTTATAC |
| HA-DDX43-F | GTTCCAGATTACGCTACTGATCGCGACGATGATGA |
| HA-R | AGCGTAATCTGGAACATCGTAT |
| DDX43-D399A-F | GTTTTAGCTGAGGCAGATAGGATGTTAGACATG |
| DDX43-D399A-R | TGCCTCAGCTAAAACAATATATGAGAAATTTATG |
| pFastBac-5'out-R | GGTTTCGGACCGAGATCCG |
| pFastBac-3'out-F | AAGCTTGTCGAGAAGTACTAG |
| pFastBac-6HFLAG-F | TCTCGGTCCGAAACCATGCATCATCATCATCATCATGATTACAAGGATGACGACGAT |
| pFastBac-Spn-E-R | TTCTCGACAAGCTTTTATGTTTGGAACATCTGAAAATC |
| SBP-Spn-E-F | GCCAGCGGGAGCCCGATGAATTAAAAGCATTCTTCAATTC |
| pcDNA5-Spn-E-R | GCATTACTTAGCTATGTTTGGAACATCTGAAAATCT |
| pCold-3'out-F | TAGGTAATCTCTGCTTAAAAGC |
| pCold-5'out-R | CCTACCTTCGATATGATGATG |
| pCold-DDX43-F | ATATCGAAGGTAGGACTGATCGCGACGATGATG |
| pCold-DDX43-R | GCAGAGATTACCTACCATCTGCCTCTCCCTCT |
| pCold-DDX6-F | ATATCGAAGGTAGGATGACCGAAAATAGAATTAGTTC |
| pCold-DDX6-R | AGCAGAGATTACCTACTTGTCGCCCAGATCTTC |
| piR484-A_s | GATCCTACCAATCGGGCGTTGCTGTTGCAATTGAAGGAATGAAGACGAAGAAAGTATTGATGAACAGCACAATA |
| piR484-A_as | AGCTTATTGTGCTGTTCATCAATACTTTCTTCGTCTTCATTCCTTCAATTGCAACAGCAACGCCCGATTGGTAG |
| pIB-BmAgo3_PiggyBac_F | TATCAATCGGGCGTTGCTGTTGCAATTCGGTCTCGATTCTACGC |
| pIB-BmAgo3_PiggyBac_R | AATTGCAACAGCAACGCCCGATTGATAAGGAGAGGGTTAGGGATAGGCTTAC |
| Target RNA preparation | |
| BmAgo3 target RNA 5' fragment | AGAGUCCUUCGAUAGGGACAAGACAAUUGCACUGAUCAUGAACUCCUCUCUUCCCGCAGACAGCAAAUUCUCA |
| BmAgo3 target RNA 3' fragment | UGCUUUUCCUUUUAUACAACCGUUCUACACUCAACGCGAUGUAAAUC |
| BmAgo3 target bridge oligo | TATAAAAGGAAAAGCATGAGAATTTGCTGTC |
| dsRNA preparation | |
| dsRluc (*Renilla luciferase*) -F | TAATACGACTCACTATAGGGCCTTTCACTACTCCTACGAGC |
| dsRluc (*Renilla luciferase*) -R | TAATACGACTCACTATAGGGTGGAGCGTCCTCCTGGCTG |
| dsSiwi-F | GCGTAATACGACTCACTATAGGATCACCCCAGAAAGACAACG |
| dsSiwi-R | GCGTAATACGACTCACTATAGGCTGTGCACGTATGGGATTTG |
| dsSpnE-CDS-F | TAATACGACTCACTATAGGGACCGCAAGATCATTCTTTCCA |
| dsSpnE-CDS-R | TAATACGACTCACTATAGGGTCCGTAGACATAGCCGAGCA |
| dsSpnE-UTR-F | TAATACGACTCACTATAGGGACCGTTTATTTTTAACATAACTG |
| dsSpnE-UTR-R | TAATACGACTCACTATAGGGGCCATATTTTCAATTTCACTTC |
| dsDDX43#1-F | TAATACGACTCACTATAGGGACCGATAAAGTAGGTAGGG |
| dsDDX43#1-R | TAATACGACTCACTATAGGGCAGCTTATCCCATCTGTTGT |
| dsDDX43#2-F | TAATACGACTCACTATAGGGACATGGCCTACTGGAGTAC |
| dsDDX43#2-R | TAATACGACTCACTATAGGGATGCAACATCAGTGGCAATC |
| dsDDX6-F | TAATACGACTCACTATAGGGCGTGTGTTTCACGACTTTCG |
| dsDDX6-R | TAATACGACTCACTATAGGGTTGGGAATTGGCTTGATTTC |
| Quantitative real-time PCR | |
| *rp49* -F | AGGTATTGACAACAGAGTCC |
| *rp49* -R | GGAGCATATGACGGGTCTTC |
| *DDX43* -F | GTAGTCGTACAGACACCGGC |
| *DDX43* -R | TATTACGGCCGCGATTTCCA |
| Probes for northern blot analysis | |
| piR484-A | TCATTCCTTCAATTGCAACA |
| Artificial_Siwi_piRNA_PiggyBac | TACGCGTAGAATCGAGACCGAATTGCAACA |
